# Translation of upstream open reading frames in a model of neuronal differentiation

**DOI:** 10.1101/412106

**Authors:** C.M. Rodriguez, S.Y. Chun, R.E. Mills, P.K. Tod

**Affiliations:** Department of Neurology, University of Michigan, Ann Arbor, MI, USA; Neuroscience Graduate Program, University of Michigan, Ann Arbor, MI, USA; Department of Computational Medicine and Bioinformatics, University of Michigan, Ann Arbor, MI, 48109, USA; Department of Human Genetics, University of Michigan, Ann Arbor, MI, 48109, USA; VA Ann Arbor Healthcare System, Ann Arbor, MI, USA

**Keywords:** Translation, Ribosome profiling, Upstream open reading frame, Near-cognate start codon, 5’ untranslated region, Neuronal differentiation

## Abstract

Upstream open reading frames (uORFs) initiate translation within mRNA 5’ leaders, and have the potential to alter main coding sequence (CDS) translation on transcripts in which they reside. Ribosome profiling (RP) studies suggest that translating ribosomes are pervasive within 5’ leaders across model systems. However, the significance of this observation remains unclear. To explore a role for uORF usage in neuronal differentiation, we performed RP on undifferentiated and differentiated human neuroblastoma cells. Using a spectral coherence algorithm (SPECtre), we identify 4,954 uORFs across 31% of all neuroblastoma transcripts. These uORFs predominantly utilize non-AUG initiation codons and exhibit translational efficiencies (TE) comparable to annotated coding regions. Usage of both AUG initiated uORFs and a conserved and consistently translated subset of non-AUG initiated uORFs correlates with repressed CDS translation. Ribosomal protein transcripts are enriched in uORFs, and select uORFs on such transcripts were validated for expression. With neuronal differentiation, we observed an overall positive correlation between translational shifts in uORF/CDS pairs. However, a subset of transcripts exhibit inverse shifts in translation of uORF/CDS pairs. These uORFs are enriched in AUG initiation sites, non-overlapping, and shorter in length.

Cumulatively, CDSs downstream of uORFs characterized by persistent translation show smaller shifts in TE with neuronal differentiation relative to CDSs without a predicted uORF, suggesting that fluctuations in CDS translation are buffered by uORF translation. In sum, this work provides insights into the dynamic relationships and potential regulatory functions of uORF/CDS pairs in a model of neuronal differentiation.

## Introduction

Alterations in protein expression and abundance are required for successful and stable cellular differentiation (Hershey, Sonenberg et al. 2012). While changes in mRNA levels provide a partial view of networks driving such cellular changes, differences in translational efficiency (TE) act as an independent contributor to this process (Brar 2016). Determining ribosomal occupancy across the transcriptome through ribosomal profiling (RP) provides is a powerful tool for assessing the relationship between mRNA abundance and translational output (Ingolia,Ghaemmaghami et al. 2009). In particular, RP in cells and organisms has revealed a detailed picture of condition-specific changes in mRNA translation rates in multiple cellular processes from meiosis to development (Ingolia, Lareau et al. 2011, Brar, Yassour et al. 2012).

The 5’ leader (traditionally referred to as the 5’ untranslated region) of mRNAs are one well-studied source of protein synthesis regulation (Kozak 1991, Calvo, Pagliarini et al. 2009, Sonenberg and Hinnebusch 2009, Hinnebusch, Ivanov et al. 2016). 5’ leaders can regulate the synthesis of the main coding sequence (CDS) product through a variety of mechanisms (Sonenberg and Hinnebusch 2009, Hinnebusch, Ivanov et al. 2016). RNA secondary structures can impede ribosomal scanning, which decreases access of assembled 40S ribosomal preinitiation complexes to CDS initiation sites. Translation can also initiate within 5’ leaders at upstream open reading frames (uORFS). In the case of uORFs that terminate after the CDS initiation site (overlapping uORFs or “oORFs”), initiation in the 5’ leader directly competes with CDS initiation for scanning 40S ribosomes and is thus predicted to be inhibitory on CDS translation. In contrast, uORFs that terminate within the 5’ leader and before the CDS initiation site (contained uORFs or “cORFs”), ribosomes can potentially reinitiate at the CDS. Thus, cORFs sometimes bypass other 5’ leader regulatory elements and can even provide stimulatory effects on CDS translation, but may be repressive as well. uORF translation can also indirectly influence CDS translation by influencing mRNA stability (Rebbapragada and Lykke-Andersen 2009) or through interactions of newly synthesized uORF protein products with the translating ribosome (Parola and Kobilka 1994, Rabadan-Diehl, Martinez et al. 2007). As such, the relationship of each uORF to the translation of its cognate CDS can be complex, making it difficult to define their specific functions and regulation across the transcriptome based on position alone.

Early ribosome profiling reports demonstrated the surprising finding that ribosome protected fragments (RPFs) are highly enriched within 5’ leader regions of mRNAs (Ingolia, Ghaemmaghami et al. 2009, Ingolia, Lareau et al. 2011, Brar, Yassour et al. 2012). Since these first reports, there have been several studies that investigated 5’ leader translation (Bazzini, Johnstone et al. 2014, Ingolia, Brar et al. 2014, Crappe, Ndah et al. 2015, Fields, Rodriguez et al. 2015, Calviello, Mukherjee et al. 2016, Chun, Rodriguez et al. 2016). These studies revealed potential roles for uORFs in circadian clock regulation, organism development, and the cell cycle (Brar, Yassour et al. 2012, Janich, Arpat et al. 2015, Johnstone, Bazzini et al. 2016, Blank, Perez et al. 2017, Fujii, Shi et al. 2017). For example, AUG initiated uORFs were detected in the transcripts of key developmental signaling proteins during murine development (Fujii, Shi et al. 2017). Homozygous deletion of an AUG initiated uORF in the 5’ leader of *PTCH1*—which encodes the major receptor for SHH signaling—disrupted differentiation of mouse embryonic stem cells into neural progenitors (Fujii, Shi et al. 2017). Interestingly, ribosome profiling at various time points throughout neuronal differentiation of human embryonic stem cells revealed shifts in 5’ leader coverage on a number of transcripts (Blair, Hockemeyer et al. 2017).

However, these data were not systematically analyzed for active translation and characterization of uORFs, and relied solely on RPF reads as a measure translation of the whole 5’ leader. Additionally, few studies to date have included non-AUG initiated uORFs in their analysis (Brar, Yassour et al. 2012, Fijalkowska, Verbruggen et al. 2017, Spealman, Naik et al. 2017) despite their potential to contribute significantly to the pool of footprints within 5’ leaders.

Treatment of human neuroblastoma cells with retinoic acid triggers their exit from the cell cycle and their differentiation into a neuron-like cell type (Sidell 1982, Pahlman, Ruusala et al. 1984). While many studies have sought to understand genetic changes underlying this process, most have focused on transcript-level changes, with evaluation of shifts in protein abundance only studied on a case-by-case basis (Hanada, Krajewski et al. 1993, Kaplan, Matsumoto et al. 1993, Korecka, van Kesteren et al. 2013, Pezzini, Bettinetti et al. 2017). Here we used RP in this simple model system to study the role of uORF activity in regulating protein translation during neuronal differentiation. Using a spectral coherence algorithm (SPECtre) and stringent dataset filtering we defined a set of translated uORFs, the majority of which initiate at a near-cognate start codon (Chun, Rodriguez et al. 2016). The presence of an AUG or conserved non-AUG uORF predicted lower CDS TE independent of cell state. Moreover, there were significant shifts in uORF usage that occurred with neuronal differentiation, suggesting a potential regulatory role. We observe less of a differentiation-dependent shift in translation of CDSs downstream of a uORF expressed across conditions, suggesting that uORFs act as a translational buffer on the transcripts in which they reside. Together this work provides important insights into how uORFs may function to regulate the translation of their associated CDS in a model of neuronal differentiation.

## Results

### Ribosome profiling detects conditionally regulated translation with differentiation

We first confirmed the efficacy of RA treatment in differentiating SH-SY5Y cells. Cells were propagated to 80% confluency prior to 10μM RA treatment for six days (**Figure 1A**). RA treatment induces an exit from the cell cycle and a change in cellular morphology. Previous studies have used a similar protocol as a model for dopaminergic neuronal differentiation, although RA treatment is thought to generate a more immature neuron-like cell than what can be achieved from a neural progenitor (Sidell 1982, Pahlman, Ruusala et al. 1984, Kaplan, Matsumoto et al. 1993). Cytoskeletal alterations confirm a shift towards a more neuron-like state after 6 days of treatment. Cytoplasmic beta-actin immunofluorescence decreased and neurofilament labeled neurites increased in length in the differentiated cells (**Figure 1B-D**) (Micheva, Vallee et al. 1998, Cheever and Ervasti 2013). We also observed an increase in expression of FMRP, a protein involved in neuronal function and translational control that is highly expressed in neurons relative to other cell types and tissues (**Figure 1E-F**) (Hinds, Ashley et al. 1993).

**Figure 1:**
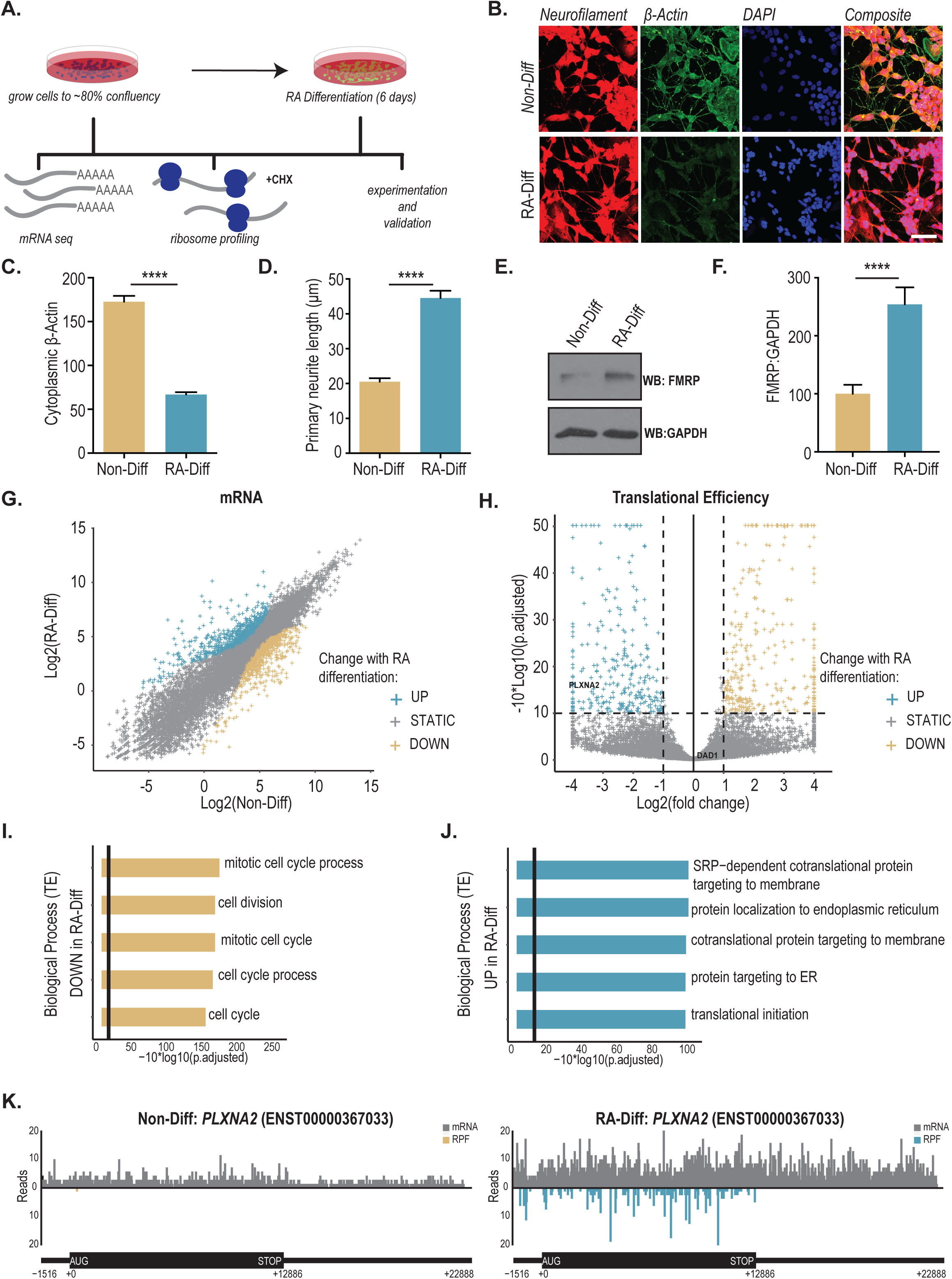
Retinoic acid treatment induces differential translation in SH-SY5Y human neuroblastoma cells. A) Schematic of experimental design and data acquisition work-flow. B) Immunocytochemistry performed on Non-Diff and RA-Diff SH-SY5Y cells with antibodies against neurofilament (red) and β-actin (green). Nuclei were DAPI stained (blue). C) β-actin expression was decreased in RA-Diff cells. Individual cell fluorescence was quantified and represented as a corrected total cell fluorescence (CTCF) for β-actin; n=119 for Non-Diff and n=118 for RA-Diff. D) Primary neurite length measured by neurofilament staining; n=109 for Non-Diff and n=100 for RA-Diff. E) FMRP expression by immunoblot before and after RA treatment, quantified in F); n=4 for both conditions. For panels C), D), and F) Student’s t test, ****p≤0.0001. Graphs represent mean ± S.E.M. G) Differential mRNA abundance based on Non-Diff versus RA-Diff TPM. Transcripts were defined as significantly up-regulated (cyan) or down-regulated (gold) in the RA-Diff condition based on rank-change in abundance compared to the Non-Diff condition. H) Volcano plot of transcripts with differential translation by translational efficiency (TE) by Riborex analysis. Significantly up-regulated genes (cyan) and down-regulated genes (gold) in RA-Diff cells are defined by an absolute log2-normalized fold-change cutoff of ±1 (vertical lines), and a multiple testing corrected p-value cutoff of 0.1 (horizontal line). I) Gene sets (biological process) with significantly downregulated TE in RA-Diff cells. Genes upregulated in RA-Diff cells is shown in (J). The top five biological processes with significant change using a multiple testing corrected p-value cutoff of 0.05 (vertical line) are shown on the graph. K) Plot shows normalized mRNA reads (grey) and RPF (cyan/gold) over the 5’leader (thin line, left), and CDS (thick line, middle). The axon guidance gene, *PLXNA2*, is representative of a transcript with higher translational efficiency and RPF in the RA-Diff condition.

Global mRNA sequencing (mRNA-Seq) was performed to determine underlying transcriptional differences in undifferentiated (“Non-Diff”) or RA Differentiated (“RA-Diff”) cells (**Figure 1G**). Alignment statistics are summarized in **Supplemental Table S1**. Gene ontology (GO) analysis revealed downregulation of a network of transcripts associated with the biological processes of mitotically active cells in the RA-Diff cells (**Supplemental Figure S1A**). In contrast, transcripts comprising biological pathways associated with cell communication and stimulus response were upregulated in this condition (**Supplemental Figure S1B**). GO analysis of transcript groups for “cellular compartment” and “molecular function” reveal similar changes associated with both conditions (**Supplemental Figure S1C-F**).

Ribosome profiling can resolve the specific regions of mRNA undergoing translation at nucleotide resolution across the transcriptome within a cell population (Ingolia, Ghaemmaghami et al. 2009). By comparing ribosomal occupancy within a given transcript in Non-Diff to RA-Diff cells, we are able to gauge translational differences. This can be accomplished by normalizing RPF abundance in the CDS to mRNA expression in samples prepared in parallel as a measure of TE (**Figure 1H**). Inspection of biological processes by GO analysis with significant translational efficiency changes revealed mainly a downregulation of transcripts encoding proteins involved in mitosis in the RA Diff cells (**Figure 1I**). Transcripts involved in endoplasmic reticulum (ER) function were significantly upregulated in the RA Diff cells (**Figure 1J**).

Investigation of transcript groups associated with a specific molecular function or cellular compartment further clarify the translational changes associated with neuronal differentiation (**Supplemental Figure S2A-D**). A more complete view of translation on an individual transcript is exemplified by *PLXNA2* which encodes a membrane-bound protein involved in nervous system development and axon guidance (Van Vactor and Lorenz 1999). Its mRNA coverage is upregulated upon differentiation; however, the increased expression is much greater at the RPF level (**Figure 1K**), producing a higher translational efficiency. Other transcripts such as *DAD1,* a factor critical for N-terminal glycosylation with roles in apoptosis and the unfolded protein response, exhibit shifts at both the mRNA and RPF level which produces no significant change in translational efficiency (**Supplemental Figure S2E**) (Kelleher and Gilmore 1997). All mRNA, RPF, and TE changes are detailed in **Supplemental Table S2**, and all GO analysis results are detailed in **Supplemental Table S3**.

### Characterization and experimental validation of SPECtre-identified uORFs

To annotate uORF sequences within the 5’ leader of mRNA, we utilized the SPECtre algorithm for classifying active regions of translation (Chun, Rodriguez et al. 2016). SPECtre accounts for the fundamental attribute of an actively translating ribosome to shift position three nucleotides at a time as it synthesizes new peptides and the ability of ribosome profiling to resolve this behavior with peaks in read coverage. Our algorithm takes an unbiased approach to scoring all potential uORFs from start site to the next in-frame stop codon (**Figure 2**). Start sites were predefined based on results from ribosome profiling studies using the drug harringtonine which halts ribosomes at the site of initiation. Thus, potential uORFs were included if they initiated at an AUG, AGG, ACG, AAG, AUC, AUA, AUU, CUG, GUG, and UUG. For each potential uORF, the pattern of read coverage within this designated sequence is compared against the pattern of reads across all known protein-coding regions in the experimental library. This analysis results in a set of experimentally determined scores that are then subjected to a range of transcript-level filters.

**Figure 2:**
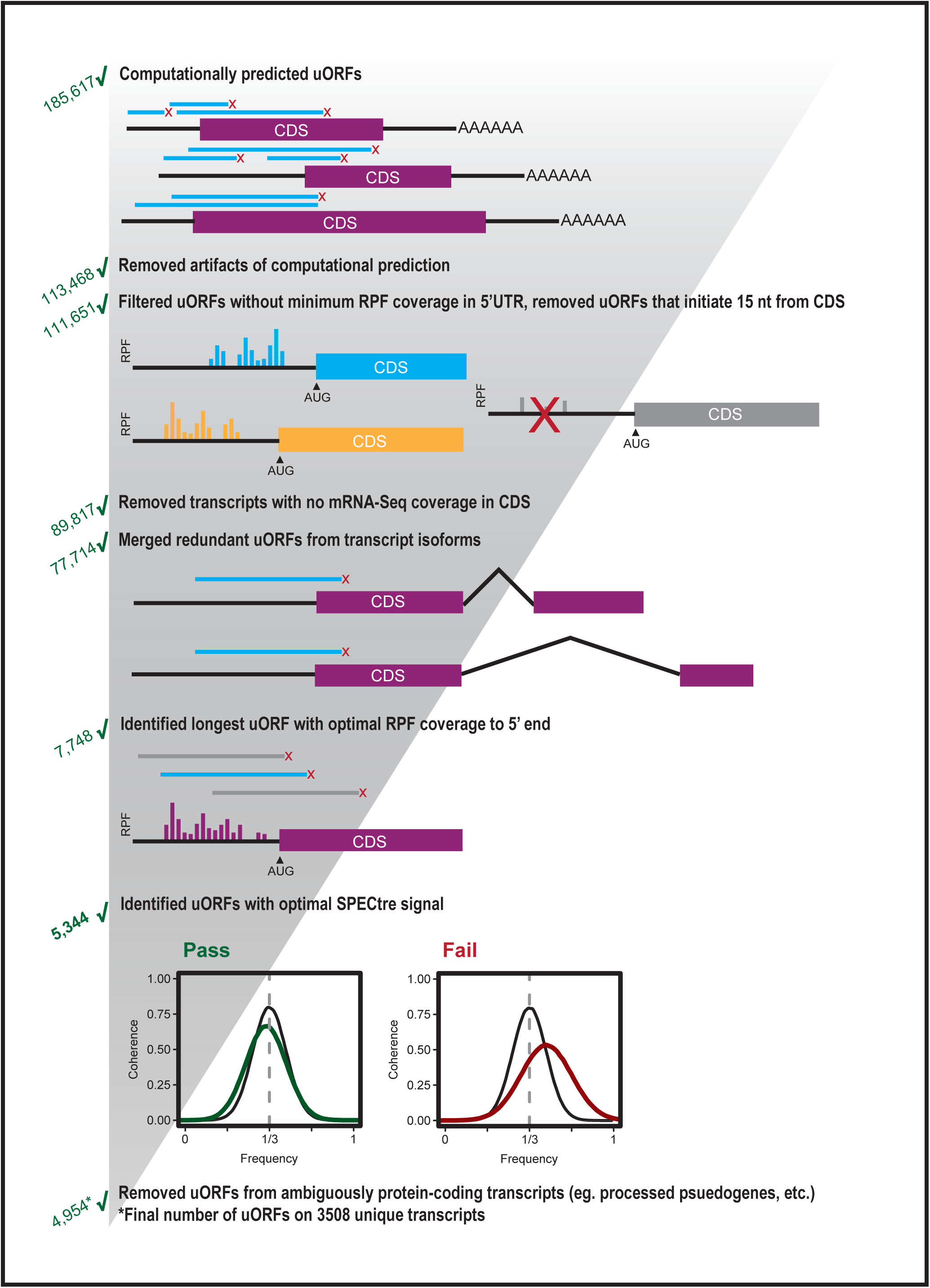
Computational prediction and filtering of upstream-initiated open-reading frames. ORFs were predicted from AUG and non-AUG, near-cognate translation initiation sites in the 5’ leader of annotated protein-coding genes, and computationally extended to the first termination site encountered in the 5’ leader (upstream-terminated ORFs) or CDS (overlapping ORFs). Predicted ORFs were then screened through a series of heuristic filters including: 1) minimum RPF coverage in the 5’ leader, 2) minimum mRNA-seq coverage in CDS, 3) in-frame N-terminal extensions, 4) redundant isoforms, 5) minimum length with optimal RPF coverage, 6) sufficient SPECtre signal, and 7) removal of ambiguously annotated protein-coding transcripts.

We established a translational threshold based on the distribution of scores in known coding genes to establish a minimum SPECtre score needed to classify a region as actively coding with a 5% false discovery rate (FDR) allowed. This results in a set of 3,508 transcripts with 4,954 unique uORFs (**Figure 2, Supplemental Table S4**). Of these transcripts, 1,599 contained overlapping upstream-initiated ORFs (specified as oORFs), 1,438 uORFs fully contained in the 5’ leader (cORFs), and 471 transcripts had two or more uORFs of either of these two categories (**Figure 3A**). The median distance of the uORF initiation site from the CDS is 99 nucleotides (**Figure 3B**). uORFs have a median length of 78 nucleotides, but can span upwards of 500 nucleotides in length (**Figure 3C**).

**Figure 3:**
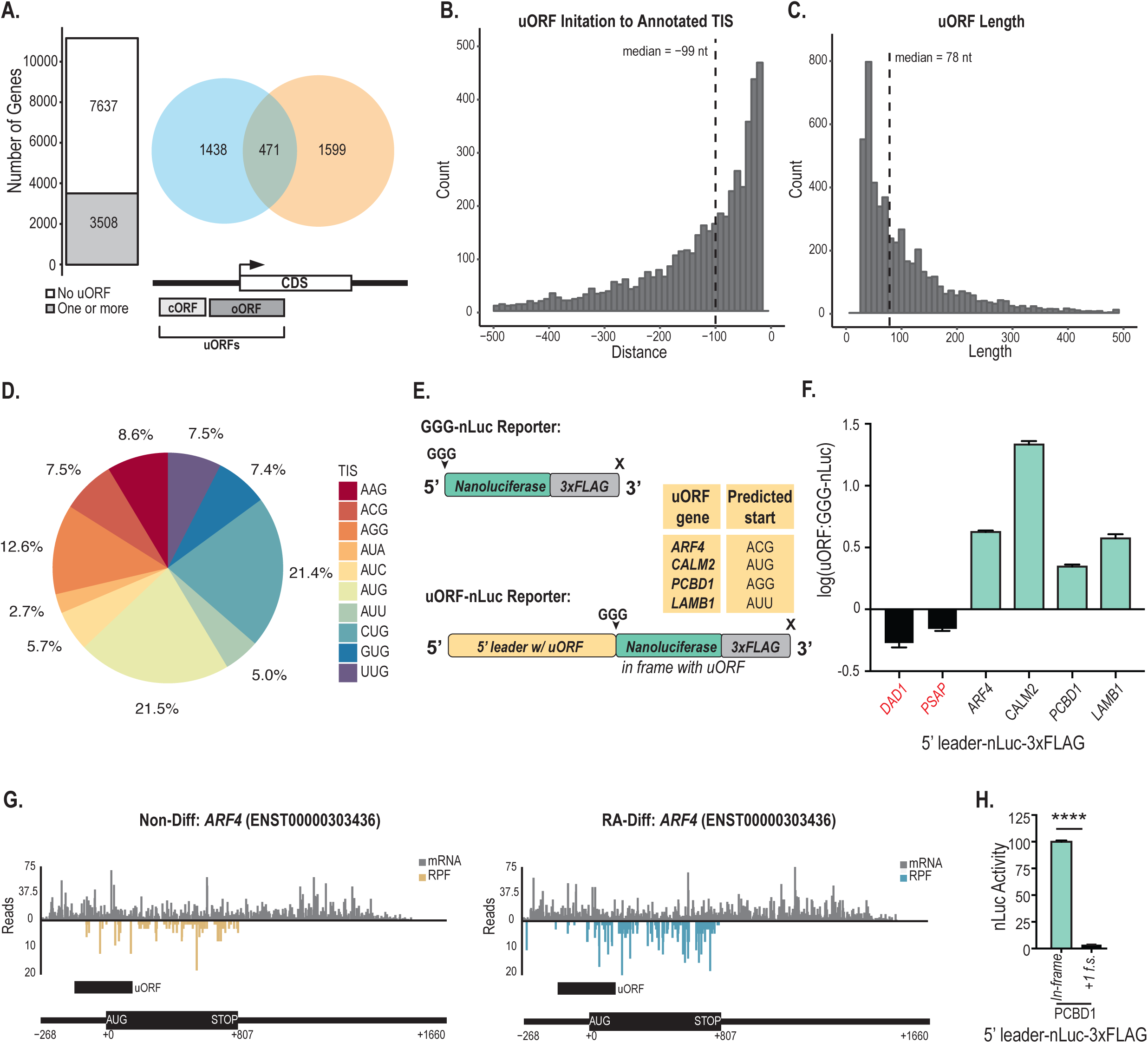
Characterization and validation of predicted uORFs. A) The number of genes with at least one predicted ORF (bar plot) in the 5’ leader of evaluated protein-coding genes. The number of genes with a predicted ORF terminated upstream in the 5’ leader only (orange), terminated in the CDS only (blue), or with both a predicted upstream- and CDS-terminated ORF (overlap). B) Distribution of predicted ORF translation initiation position relative to the annotated protein-coding CDS start site. C) Distribution of predicted ORF lengths. D) Distribution of uORF translation start sites (TIS). AUG represents all AUGs predicted by SPECtre, or upstream/downstream 30-nt from the SPECtre predicted start site if no intervening stop codon is present. Near-cognate start codons are utilized in the majority of uORFs, while AUG is the single most common start site. E) Schematic of the uORF nanoluciferase (nLuc) reporters used in this study. GGG-nLuc serves as a negative control, as its AUG initiation start codon is mutated to a GGG codon. This reporter supports very little translation. A table of the predicted start sites for each uORF reporter. F) nLuc assays performed in SH-SY5Y cells confirmed expression of these uORFs (teal). 5’ leaders not included in the uORF dataset (black) are below the GGG-nLuc reporter activity and considered not translated. All values are normalized to the GGG-nLuc control performed in parallel, data for individual reporters was collected in triplicate in multiple experiments. Student’s t test, all teal uORFs in panel F) have a p value ≤0.0001. Graph represents mean ± S.E.M. G) Plots show mRNA reads (grey) and RPF counts (cyan/gold) for *ARF4*. The annotated uORF is characterized by the presence of consistent RPF coverage in the 5’ leader. H) Frameshifting the uORF relative to nLuc decreases translation of the reporter. The reporter was cloned so that the nLuc tag was frameshifted (f.s.) out of frame with the predicted uORF and the CDS start site. n=3, Student’s t test, ****p≤0.0001. Graph represents mean ± S.E.M.

Previous work using harringtonine, a drug that stalls ribosomes at initiation sites, revealed a surprising occurrence of near-AUG codons enriched in ribosome peaks (Ingolia, Lareau et al. 2011). Though near-cognate initiation had been recognized previously, this hinted that there may be a greater number of initiation events at these codons than previously expected (Zitomer, Walthall et al. 1984, Peabody 1987, Kozak 1989, Peabody 1989, Mehdi, Ono et al. 1990, Ingolia, Ghaemmaghami et al. 2009). When inspecting the translation start site of each SPECtre-identified uORF, we see a number of non-canonical initiation events.

Translation start sites were plotted to show the relative contribution of each in the final dataset (**Figure 3D**). It is important to note that this is the breakdown of start sites within the constraint of our initial parameters, which limit potential start codons to those with a preset identity. AUG initiation sites were accounted for by two different methods: they were either directly identified by SPECtre, or factored into the total count if they were present within 30 nucleotides upstream or downstream of the start of the SPECtre signal without an intervening stop site. Due to the high potential translatability of ORFs with AUG start codons, these were all annotated as AUG start sites. This constitutes 21.5% of the initiation sites used. In comparison, we detected 21.4% of uORFs use CUG as their initiation codon, consistent with previous reports (Peabody 1987, Ingolia, Lareau et al. 2011).

We next looked to validate our findings using orthogonal techniques. A straight forward approach would be to use mass spectrometry to detect uORF derived peptides; however this has previously been proven difficult, likely due to a range of complications from sample preparation, bias in annotation algorithms, as well as intrinsic factors that make these peptides difficult to detect (Bazzini, Johnstone et al. 2014, Chugunova, Navalayeu et al. 2018). Several methods have been developed to enhance the detection of small peptides, each with variable yields (Slavoff, Mitchell et al. 2013, Bazzini, Johnstone et al. 2014, Na, Barbhuiya et al. 2018). We therefore measured active translation of uORFs in our dataset by creating nanoluciferase (nLuc) reporters. For each candidate evaluated, the complete 5’ leader upstream of the start site through the entire predicted coding region of the uORF was placed upstream of an nLuc tag without an AUG start codon (**Figure 3E**). GGG-nLuc alone was used as a negative control (Kearse, Green et al. 2016). Using this system, we confirmed SPECtre identified uORFs residing in the 5’ leader of 4 genes: *ARF4*, *CALM2*, *PCBD1*, and *LAMB1*. *ARF4*, *PCBD1,* and *LAMB1* are predicted to utilize near-cognate start sites, while *CALM2* utilizes an AUG (**Figure 3E**). Reporters were co-transfected into SH-SY5Y cells with pGL4.13 which encodes firefly luciferase (FFluc) as a transfection control. *DAD1* and *PSAP* serve as negative controls, as their 5’ leaders were filtered out early on in our analysis. All 4 of our predicted uORFs showed a significant level of translation above GGG-nLuc (**Figure 3F**). RPF and mRNA coverage of *ARF4* reveals significant coverage across the uORF in the Non-Diff state (**Figure 3G**). One key feature of SPECtre is its ability to discriminate reading frame (Chun, Rodriguez et al. 2016).

While we have shown that 5’ leaders can be translated, we wanted to investigate whether genes identified by SPECtre supported spurious translation or if these sequences were specific to initiation in one reading frame. To evaluate this, we mutated the reporter for *PCBD1* so that the predicted uORF was out of frame. Placing nLuc out of frame resulted in a significant drop in signal (**Figure 3H**), revealing that the ribosome discerns coding regions based on specific sequence context, and that a 5’ leader can support a discrete translation event.

### uORF/CDS pairs exhibit positively correlated TE shifts with differentiation

We next sought to determine whether SH-SY5Y differentiation can affect uORF SPECtre score. We performed k-means clustering using the SPECtre score of uORFs in the Non-Diff and RA-Diff datasets (**Figure 4A**). This revealed that a significant fraction of uORFs exhibit a high degree of cell-state specificity (blue and gold) compared to uORFs with consistent scores in both states (grey). A similar relationship was observed for uORF TE (**Supplemental Figure S3A**). These findings suggest that usage of specific uORFs may be regulated during neuronal differentiation. If these shifts in uORFs occur mostly as a means of regulating CDS translation, then we would predict an inverse relationship between these shifts in uORF usage and the TE of their cognate CDS. However, these clusters were not predictive of an inverse directional change in CDS TE (**Figure 4B**). Instead, we observed a positive correlation between CDS TE and uORF TE in this dataset (**Supplemental Figure S3B**) that was present regardless of whether we considered all uORF, or cORFs and oORFs individually (**Figure 4C-D**). These findings are consistent with a number of previous RP studies (Brar, Yassour et al. 2012, Chew, Pauli et al. 2016) and have generally been interpreted as evidence for the effects of enhanced pre-initiation complex binding and stochastic leaky scanning. While this positive relationship does not preclude the potential for specific uORFs to act as repressors, it suggests that global shifts in uORF usage are not driving the majority of changes in CDS TE observed with neuronal differentiation.

**Figure 4:**
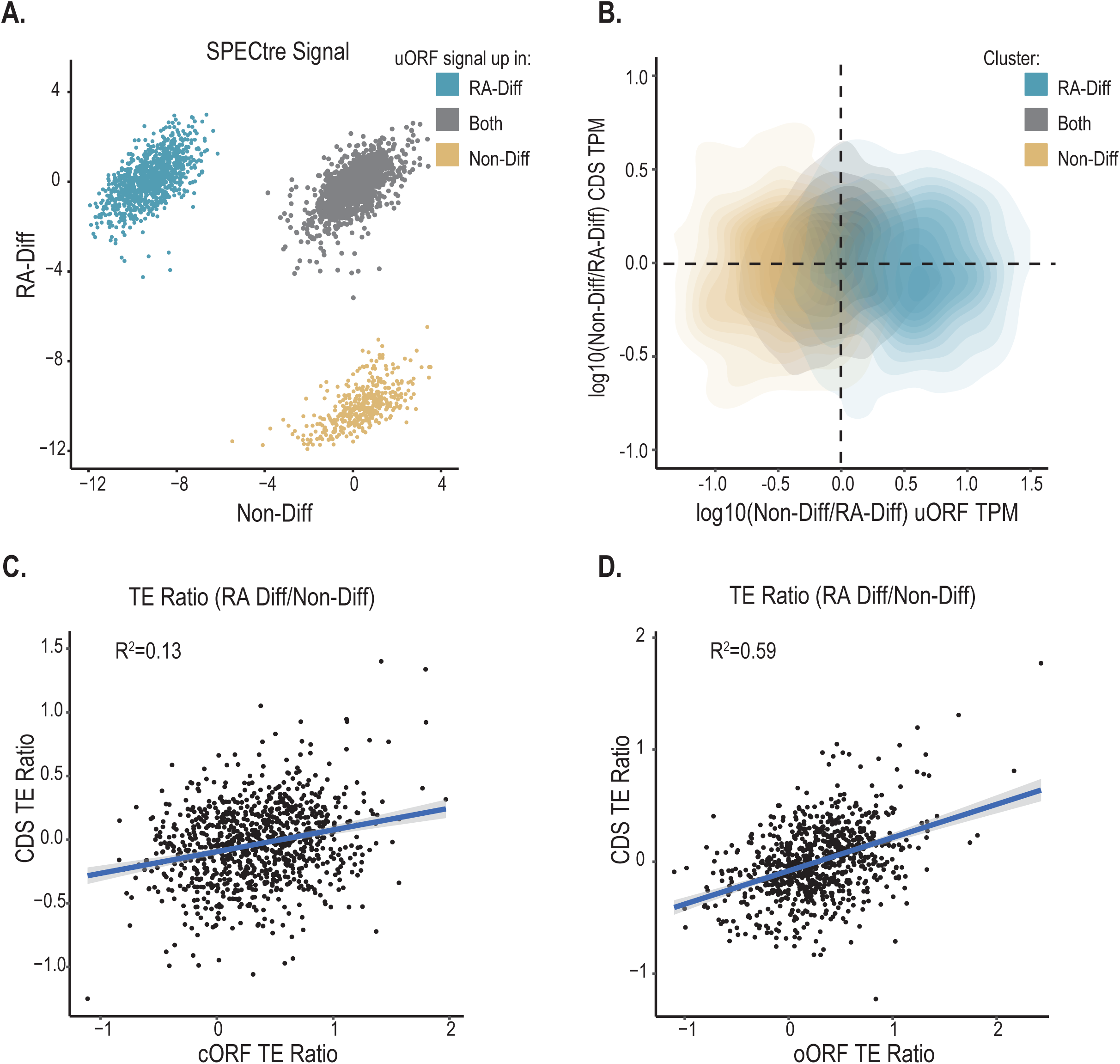
uORFs shift translationally with neuronal differentiation A) K-means clustering analysis of log2(uORF SPECtre Score) in Non-Diff and RA-Diff cells, reveals differentiation-associated shifts. Three clusters of uORF translation emerge: those that are up-regulated in RA-Diff cells (cyan), up-regulated in Non-Diff cells (gold), and uORFs with no change in translational potential (gray). B) Clustering in (A) does not correlate with directional CDS changes. Kernel density estimation analysis of changes in TPM over annotated protein-coding CDS as a function of changes in TPM over predicted upstream-initiated ORFs. Cluster identity of predicted ORF changed in translational potential as scored by SPECtre predicted ORFs enriched for translation in RA-Diff cells (cyan), predicted ORFs with enriched translation in Non-Diff cells (gold), and those with static translation across the two conditions (black) are annotated to protein-coding CDS with higher RPF abundance in Non-Diff cells (above horizontal line), and those with higher RPF abundance in RA-Diff cells (below horizontal line). C) Analysis of transcripts with a cORF reveals a positive correlation of cORF TE and CDS TE. Pearson correlation, R^2^=0.13. D) The same is true for oORFs with a Pearson correlation, R^2^=0.59.

### Robustly expressed and conserved uORFs exhibit greater inhibition of CDS translation

To further explore the relationship between uORFs and CDS translation, we limited our analyses to a stricter dataset with uORFs robustly identified in multiple technical assays in order to parse out the relationship between these two regions. We reasoned that our inclusion of non-AUG initiated uORFs, which in principle have lower potential for translation, might contribute a higher level of variability to our analysis. After filtering down to this stringent dataset, we obtained 158 overlapping ORFs (oORFs) and 137 constrained ORFs (cORFs) that initiate at either an AUG or near-cognate codon (**Figure 5A**). These cORFs and oORFs have comparable translational efficiencies (**Figure 5B**).

**Figure 5:**
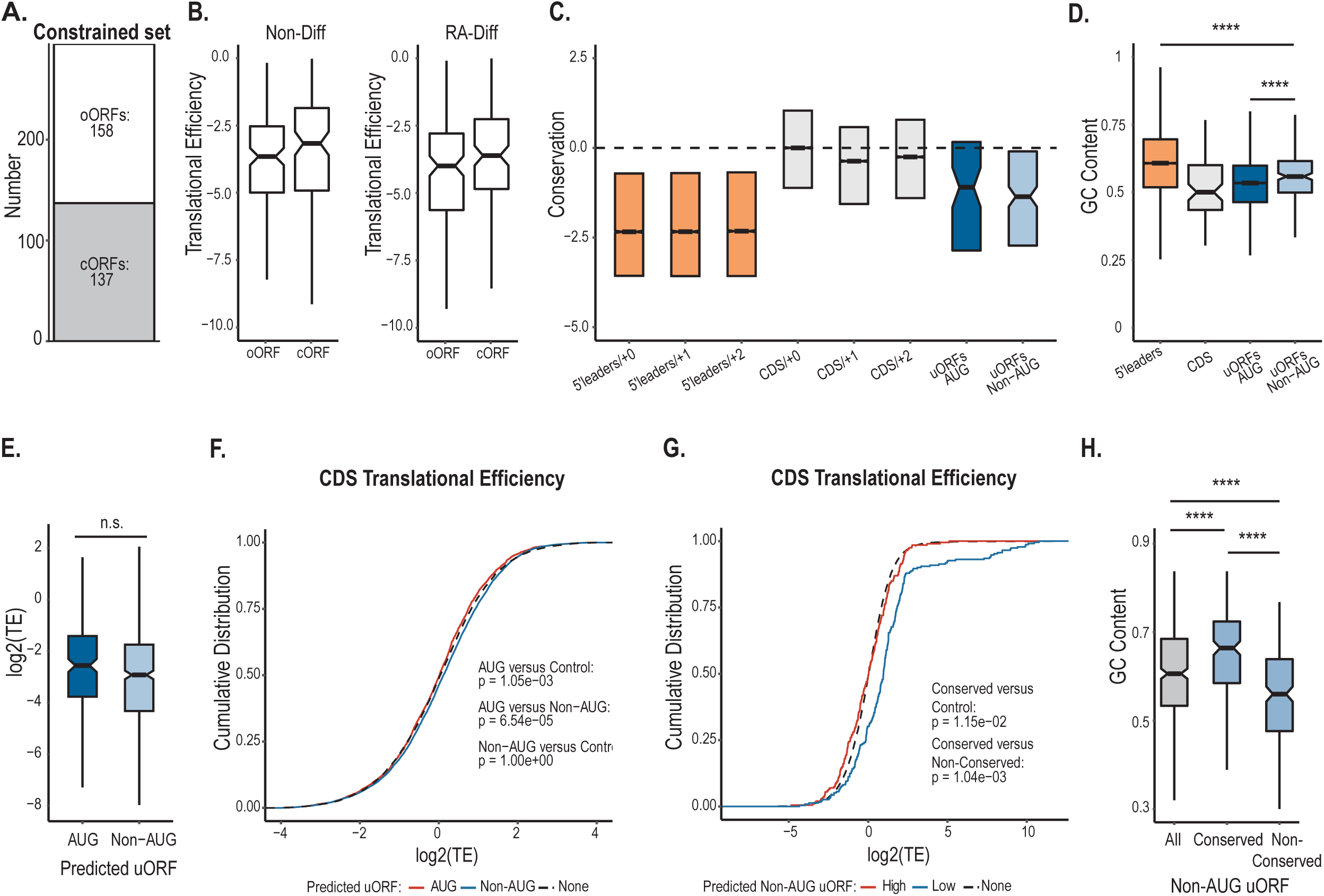
Constrained analysis of the uORF dataset reveals a repressive effect of highly conserved uORFs. A) SPECtre identified uORFs were filtered to include only uORFs that have coverage in all four Non-Diff and RA-Diff libraries; these are considered highly translated. B) Average TE values for cORFs and oORFs in the Non-Diff (left) and RA-Diff (right) conditions. C) Conservation analysis of annotated 5’ leaders in all three reading frames (orange), annotated CDS regions over all three frames (grey), predicted AUG-initiated uORFs (dark blue), and predicted non-AUG uORFs (light blue). D) Average GC nucleotide content is shown for 5’ leader regions (orange), CDSs (grey), AUG uORFs (dark blue), and non-AUG uORFs (light blue). For oORFs, only the 5’ leader region of the oORF is included. 5’ leaders are significantly more GC rich than both AUG uORFs and non-AUG uORFs, p= 5.72e-12 and 1.54e-07, respectively. Non-AUG uORFs are significantly more GC rich than CDSs and AUG uORFs, p=7.92e-18 and 2.16e-06. E) Average TE for AUG uORFs and non-AUG uORFs reveals no difference between the two subtypes. F) Empirical cumulative distribution of TE in all CDSs (black) versus CDSs from transcripts with two subsets of uORFs: those with an AUG initiation site (red) and those with a non-AUG initiation site (Blue). G) Empirical cumulative distribution of TE in all CDSs (black) versus CDSs from transcripts with two subsets of non-AUG uORFs: those in the highest quartile of conservation (Conserved, red) and those in the lowest quartile of conservation (Non-Conserved, Blue). Distributions are significantly different with p-values annotated on graph. H) GC content of non-AUG uORFs grouped by conservation. All versus conserved: p=5.89e-08, all versus non-conserved: p=2.54e-06, conserved versus non-conserved: p=1.62e-18.

We next analyzed the conservation of this set of uORFs divided into subsets based on initiation codon. uORFs were analyzed at the codon level (comprised of only the 5’ leader region for oORFs) compared to all 5’ leader sequences and annotated CDSs. Both non-AUG and AUG uORFs exhibit a range of conservation, with a significant portion showing conservation levels that are comparable to CDSs (**Figure 5C**), and enhanced conservation compared to all 5’ leader sequences. The GC content is generally higher in 5’ leaders than CDSs and can serve as a proxy for increased RNA structure due to the enhanced strength of GC hydrogen bonding compared to AU pairs (Banerjee 1980). Interestingly, we observe a decrease in GC content in both AUG and non-AUG uORFs compared to all 5’ leaders (**Figure 5D**). This shift brings these sequences closer to what is observed across all CDSs, although they remain significantly more GC rich than this population (**Figure 5D**) (Kozak 1980, Kozak 1986). In addition, there is a significant difference in the GC content of non-AUG uORFs relative to AUG uORFs, with more GC content associated with non-AUG mediated initiation. This could suggest that greater secondary structure is needed to enhance near-cognate start site initiation compared to AUG initiated translation, a finding that is consistent with biochemical studies of sub-optimal start site selection due to stalling of scanning ribosomes (Kozak 1990). This relationship also suggests that AUG initiated uORFs may be more functionally similar to classic CDS translation than non-AUG uORFs.

To investigate the impact of uORFs on downstream translation, we compared the cumulative distribution of CDSs without a predicted uORF to CDSs in our transcripts of interest. First, we stratified uORFs into two groups based on whether they were AUG initiated or non-AUG initiated. Despite differences in initiation sites, these two groups have similar translational efficiencies (**Figure 5E**). AUG initiated uORFs are associated with less translation from their cognate CDSs, although this effect was modest (**Figure 5F**) (Calvo, Pagliarini et al. 2009, Chew, Pauli et al. 2016, Johnstone, Bazzini et al. 2016, Spealman, Naik et al. 2017). In contrast, CDSs downstream of non-AUG initiated uORFs showed a comparable distribution to the control set. As the role of non-AUG uORFs has not been well elucidated, we wondered if there was an attribute of non-AUG uORFs that could delineate their impact on CDS translation. We reasoned that uORFs with regulatory potential would be more likely to be conserved across phylogeny. Therefore, we divided the non-AUG uORFs into two groups: those in the top quartile of conservation (conserved) and those in the bottom quartile of conservation (non-conserved). For the top quartile of conserved transcripts, the presence of non-AUG uORFs was significantly associated with CDS repression (**Figure 5G**). In contrast, non-AUG initiated uORFs in the lowest quartile of conservation were associated with highly translated CDSs. This is not due to TE levels of the conserved versus non-conserved, non-AUG uORFs (**Figure 5E**). Interestingly, the conserved group exhibited higher GC content, consistent with the finding that 5’ leader secondary structure is a predictor of CDS repression and that these features are conserved (**Figure 5H**) (Pelletier and Sonenberg 1985, Chew, Pauli et al. 2016). Thus, both AUG and non-AUG initiated uORFs can inhibit downstream translation.

### Ribosomal transcripts are enriched in the uORF dataset

Interestingly, there is a significant enrichment for ribosomal protein transcripts in the constrained set of uORFs, with 19 bearing oORFs and 4 bearing cORFs (**Figure 6A, Supplemental Figure S4**). Most uORF/CDS pairs in this group of transcripts exhibit a positive correlation in the directionality of changes in translational efficiency in response to differentiation, however the oORF on *RPS24* shows an inverse relationship with translation of the CDS (**Supplemental Figure S4**). We validated the expression of two uORFs from ribosomal protein transcripts: *RPS8* and *RPS18* by nanoluciferase assay (**Figure 6B**). These show robust reporter expression, which was validated by immunoblotting (**Figure 6C**). In 20 of the 23 ribosomal transcripts, the CDS is more highly translated in the Non-Diff state. A decrease in translation related transcripts was previously detailed in neuronal differentiation and attributed to a decrease in mTORC1 activity (Blair, Hockemeyer et al. 2017). Most ribosomal transcripts, including all transcripts in this subset, have a 5’ terminal oligopyrimidine (5’ TOP) motif which implements translational control mediated by stress and the mTOR signaling pathway (Meyuhas and Kahan 2015). uORFs on the ribosomal transcripts in our constrained dataset are either initiated downstream of the 5’ TOP motif or upstream of this motif leading to its inclusion in the uORF. The extent to which the 5’ TOP motif dictates uORF translation is unknown. However, the enrichment in uORFs in these transcripts suggests a specific role for these elements in modulating state-dependent expression of ribosomal proteins.

**Figure 6:**
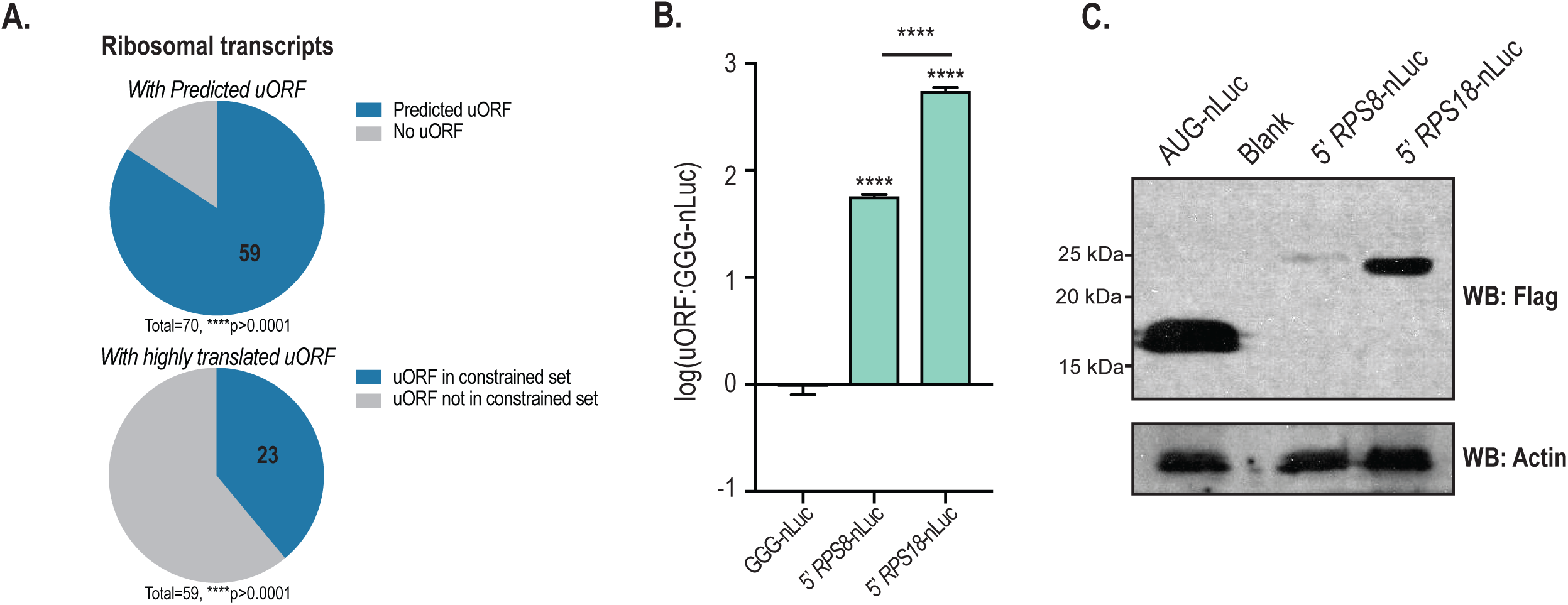
Ribosomal transcripts are enriched in the uORF dataset A) Top: Chart shows the percent of total actively translated ribosomal protein transcripts (n=70) with a predicted uORF (n=59). Bottom: Chart shows the percent of ribosomal transcripts with uORFs that are in the constrained (highly translated) dataset (n=23). Ribosomal transcripts are enriched in both sets by Fisher’s exact test, ****p>0.0001. B) nLuc assay in SH-SY5Y cells transfected with uORF reporters for ribosomal transcripts: *RPS8* and *RPS18*. C) Immunoblot of these reporters show an increase in molecular weight relative to AUG-nLuc control, confirming translation initiation within the 5’ leader of these ribosomal transcripts.

### uORFs in the constrained dataset buffer against differentiation-induced shifts in CDS TE

Given their negative impact on CDS translation (**Figure 5F-G**), we revisited our evaluation of the relationship between uORF translation and CDS translation with RA-induced cellular differentiation using our constrained dataset. Among transcripts in the constrained set, we again observed a weakly positive correlation between conditional translation of uORFs (cORFs and oORFs) and cognate CDS translation between the two conditions (RA-Diff:Non-Diff) with regression coefficients of 0.08 and 0.18 for cORF and oORF transcripts (**Figure 7A-B**). A complete list of TE changes with differentiation (TE RA-Diff/TE Non-Diff) in uORF/CDS pairs is shown in **Supplemental Figure S4**. While many pairs show similar shifts in TE with neuronal differentiation, a subset of uORF/CDS pairs exhibit inverse relationships between uORF and CDS TE across cell states. We examined the 118 transcripts that exhibit this inverse relationship for defining characteristics. We observed that there was a greater proportion of cORFs in the inverse group than oORFs (**Figure 7C**), and these are shorter in length (**Figure 7D**). Moreover, a greater percentage of uORFs with this inverse relationship to their cognate CDS translation had a predicted AUG initiation site and a higher average TE (**Figure 7E-F**). The preference for cORF and shorter translation lengths in this context may thus reflect the fact that while half of human 5’ leaders contain an AUG, the overwhelming majority of these initiate a cORF (Calvo, Pagliarini et al. 2009).

**Figure 7:**
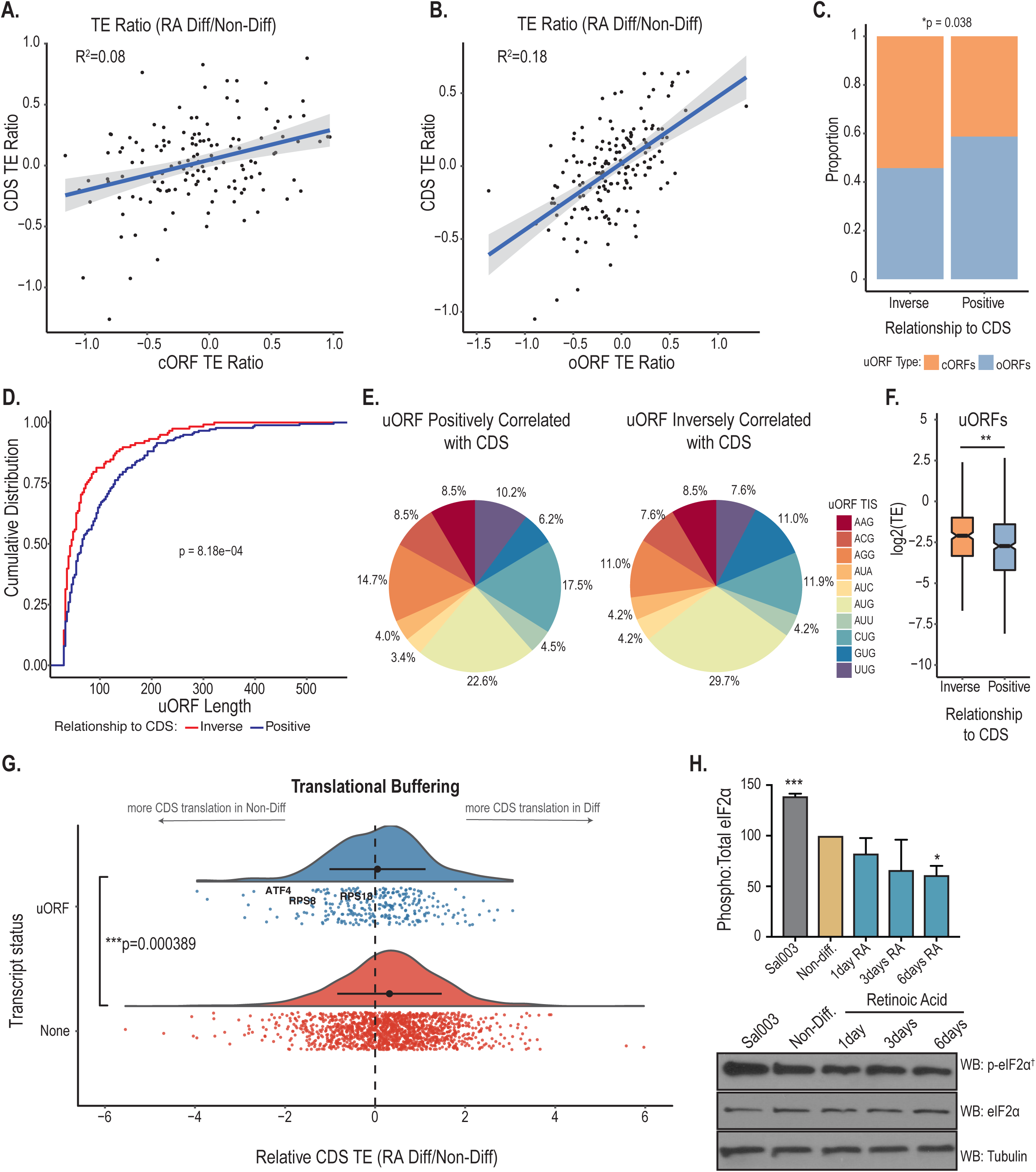
uORFs buffer against differentiation-induced shifts in CDS TE A) Analysis of the relationship between cORF and CDS translation (log10TE Non-Diff/RA-Diff) for the constrained dataset reveals that the translational efficiency of these two regions positively correlate in response to RA-Differentiation, R^2^=0.08. Regression coefficients were calculated from untransformed TE ratios. B) This was also seen for oORFs, R^2^=0.18. C-E) 40% (118/295, see Supplemental Figure S5) of uORF/CDS pairs in the constrained dataset exhibit TE shifts in the opposite direction for the uORF and the CDS with differentiation (“inverse”), while 60% of uORF/CDS pairs exhibit TE shifts in the same direction with differentiation (“positive”). C) Relative proportion of “positive” or “inverse” oORFs and cORFs. Chi-square, p=0.038. D) Distribution of uORFs by length. E) Start site codon distribution for “positive” or “inverse” uORFs. F) uORFs with an “inverse” relationship to their associated CDS have a higher average TE than those with a “positive” relationship, p=0.00923. G) Histograms of log2(CDS TE, RA-Diff/TE Non-Diff) for transcripts with a uORF in the constrained set (uORF) or no uORF (none). ANOVA, p=0.000389. H) eIF2α phosphorylation decreases after 6 days of RA differentiation. This is quantified as a relative ratio of phosphorylated to total eIF2α. † Higher sensitivity detection methods were used for phosphorylated-eIF2α, thus the absolute ratio of phosphorylated to total eIF2α is not accurately pictured.

It is worth noting that the presence of a persistently translated uORF alone can limit the translation of a CDS in a condition-dependent manner through accompanying changes in translation machinery (Andreev, O’Connor et al. 2015, Sidrauski, McGeachy et al. 2015, Starck, Tsai et al. 2016). Thus, the notion that a decrease in uORF translation is necessary to de-repress CDS translation is an incomplete view of how uORFs can function, and analysis of translation factors that may be acting *in trans* is needed to clarify the translational dynamics on individual transcripts (Young and Wek 2016). Thus we analyzed the distribution and relative TE ratio (RA-Diff/Non-Diff) on transcripts with no uORF or with a uORF in the constrained set. The range of TE ratios had a significantly increased variance in transcripts without uORFs (**Figure 7G**). These data support the role for uORFs in limiting conditional fluctuations in CDS translation (Wek 2018), while highlighting neuronal differentiation as a cellular process in which uORFs may serve their role. Transcripts without a uORF also showed a significantly lower TE ratio relative to those with a uORF in the constrained set, suggesting a larger increase in relative TE with differentiation in those transcripts (**Figure 7G**). A similar but less pronounced decrease in this TE ratio was observed for transcripts in the full set of uORFs (**Supplemental Figure S5**).

To investigate this shift in translational efficiency further, we probed for phosphorylated eIF2α, as increased eIF2α phosphorylation is known to decrease global protein synthesis (Sonenberg and Hinnebusch 2009). eIF2 must be bound to GTP (in its nonhydrolyzed form) to bind to the Met-tRNA_i_ ^Met^ initiator tRNA, this ternary complex is responsible for bringing the first amino acid to the site of translation initiation (Sonenberg and Hinnebusch 2009). Phosphorylation of the α subunit limits ternary complexes, as it acts as a competitive inhibitor of the guanine exchange factor that acts on eIF2 (eIF2B). The role of eIF2α phosphorylation in controlling global protein synthesis has largely been studied during the integrated stress response, where cell stress converges on multiple kinases that phosphorylate eIF2α (Sonenberg and Hinnebusch 2009). We detected a significant decrease in the ratio of phosphorylated eIF2α to total eIF2α after 6 days of differentiation (**Figure 7H**). While a change in eIF2α phosphorylation has not been previously described in SH-SY5Y cell differentiation, RA treatment protects cells against endoplasmic reticulum stress, and RA differentiated SH-SY5Y cells have been shown to differ from non-differentiated cells in their buffering capacity against oxidative stress inducing agents (Almeida, Vieira et al. 2017, Nair, Das et al. 2017). Further, *ATF4* is present in our list of constrained uORFs and its TE is increased in the nondifferentiated condition, consistent with past findings that *ATF4* has two uORFs whose persistent expression is key to the increase in ATF4 protein during the integrated stress response (Lu, Harding et al. 2004, Vattem and Wek 2004, Starck, Tsai et al. 2016). Taken together, this cell-state dependent shift in eIF2α may be responsible for the average increase in TE on transcripts without a uORF in the RA-Diff condition, and transcripts with uORFs may be buffered from this differentiation-dependent effect.

## Discussion

In this study, we show that application of a spectral coherence algorithm (SPECtre) to RP data and stringent filtering allows for categorization of high confidence uORFs translated in human neuroblastoma cells. The majority of identified uORFs utilize near-cognate start sites, are highly translated, and a subset of these were validated using reporter assays. As others have described, AUG initiated uORFs are associated with a repressed CDS (Chew, Pauli et al. 2016, Johnstone, Bazzini et al. 2016, Spealman, Naik et al. 2017). However, we find a similar relationship occurs with conserved non-AUG uORFs, suggesting functionality. By constraining our dataset to uORFs translated across both conditions, we further parsed out a role for uORFs in regulating protein synthesis changes with neuronal differentiation. We observe 118 uORF/CDS pairs that shift TE in opposite directions with differentiation. This aligns with the classic model of uORF usage where upregulation of uORF translation leads to a concomitant decrease in translation of the downstream CDS (Hinnebusch 1997, Wek 2018). Yet, for the majority of uORF events, this predicted inverse relationship is not observed (Andreev, O’Connor et al. 2015, Hinnebusch, Ivanov et al. 2016, Starck, Tsai et al. 2016). Instead, our data suggests that these uORFs act as buffers upon state-dependent changes in CDS translation. Changes in CDS TE downstream of a uORF were less variable in aggregate than CDSs lacking a uORF, and only CDSs lacking a uORF exhibited enhanced TE with differentiation. Together, these data support a model whereby uORFs can act as both direct regulators of downstream CDS translation and as homeostatic buffers that constrain changes in translation rates associated with cell state changes.

In yeast, near-cognate initiation in 5’ leaders is positively correlated with CDS translation, showing no apparent repressive role or repression in only a subset of transcripts (Brar, Yassour et al. 2012, Spealman, Naik et al. 2017). However, one RP study did suggest a repressive role for non-AUG uORFs under conditions of decreased start site stringency. This study utilized a knockdown of eIF1, the “gatekeeper” of start-site selection, which favors AUG initiation (Valasek, Nielsen et al. 2004, Nanda, Cheung et al. 2009). RP of eIF1 knockdown in human epithelial cells showed an increase in translation of non-AUG uORFs relative to their CDSs, but only of uORF/CDS pairs that were regulated by the knockdown (Fijalkowska, Verbruggen et al. 2017). Our data extend on this by describing that near-cognate initiation may have a role under normal cellular conditions, where endogenous translation machinery has not been manipulated. Stratifying our analysis by conservation was key to identifying this result. We show that conserved non-AUG uORFs are mildly repressive in aggregate; we suspect that using SPECtre and strict criteria to identify robust translation helped to clarify this finding. This is consistent with findings that features of AUG-initiated uORF-mediated translation are conserved (Chew, Pauli et al. 2016).

We have relied on past biochemical studies to inform our hypothesis that uORFs that do not allow for reinitiation after translation must be bypassed in order to de-repress the expression of the downstream CDS, as has been observed on *GCN4* in yeast in a process termed “delayed reinitiation” (Mueller and Hinnebusch 1986, Abastado, Miller et al. 1991, Hinnebusch 1997, Vattem and Wek 2004, Kozel, Thompson et al. 2016). There are two determining cORFs upstream of the *GCN4* CDS with the first showing persistent translation, and the second acting under normal conditions to preclude the scanning ribosome from reaching the CDS with a ternary complex for initiation (Hinnebusch 1997). In delayed reinitiation, under conditions of starvation, the second uORF is bypassed due to limiting ternary complexes which allows scanning ribosomes to continue on and load ternary complexes prior to reaching the CDS (Sonenberg and Hinnebusch 2009). Hence, GCN4 protein is upregulated.

However, several reports have suggested a positive relationship between uORF translation and CDS translation, explaining this by the low frequency of uORF translation and a reliance on whole transcript activity to increase in order to detect such a rare event (Brar, Yassour et al. 2012, Jang, Lahens et al. 2015, Chew, Pauli et al. 2016). The scanning model of translation initiation predicts that if uORF usage at suboptimal codons is an inefficient event, that it would increase as the number of preinitiation complexes increase on a transcript (Mehdi, Ono et al. 1990, Hinnebusch, Ivanov et al. 2016). Both the a priori filtering and the additional constraining of our dataset to uORFs readily detected in both conditions likely eliminated some of the noise in our analysis, however we still observe a positive correlation of shifts in translational efficiency (TE RA-Diff/TE Non-Diff) for both oORF/CDS and cORF/CDS pairs.

The positive regression coefficient for the oORF/CDS pairs illustrates that even in the setting of an overlapping uORF, which precludes reinitiation at the CDS, the positive relationship is maintained. However, for both the full uORF set and the constrained uORF set, the regression coefficient for the oORF-bearing transcripts was more positive than transcripts with cORFs. The reason for this was clarified when we investigated the characteristics of uORFs that shift opposite to their downstream CDS. The majority of these were, in fact, cORFs and more commonly utilized an AUG initiation codon. This finding is consistent with the fact that the majority of AUG codons in the 5’ leader do not produce an overlapping ORF (Calvo, Pagliarini et al. 2009). Moreover, there is a strict balance of translation initiation factors required for proper initiation at the main AUG of the CDS, thus an increase in upstream initiation at a stronger start site like an AUG is likely to decrease further scanning (Calvo, Pagliarini et al. 2009).

Earlier studies performed ribosome profiling on cells in response to robust stressors such as arsenite and tunicamycin, and showed that CDSs on transcripts with uORFs either increase in TE or remained resistant relative to the global decrease in translation (Andreev, O’Connor et al. 2015, Sidrauski, McGeachy et al. 2015). This included transcripts with known uORFs as well as transcripts that were later analyzed for uORF translation. Unexpectedly, we observed an increase in CDS TE on transcripts without a uORF in the differentiated condition relative to transcripts with a uORF in the constrained dataset. Our analysis was performed on all transcripts with persistently translated uORFs in aggregate without *a priori* filtering for the translational response of the associated CDS, which makes a strong case for an overarching role of uORFs as buffers to their associated CDS. Further, this led to the confirmation of decreased eIF2α phosphorylation in differentiated cells. We believe that this is indicative of a neuronal differentiation-dependent shift in eIF2α phosphorylation separate from an alleviation of stress, possibly associated with retinoic acid treatment itself (Roffe, Hajj et al. 2013, Almeida, Vieira et al. 2017, Nair, Das et al. 2017). Thus, our data suggests that the presence of persistently translated uORFs can delay scanning ribosomes from reaching the CDS and potentially provide more time for them to acquire a ternary complex (Andreev, O’Connor et al. 2015, Starck, Tsai et al. 2016, Wek 2018). However, the global analysis performed here is not sufficient to answer this question on individual transcripts.

One set of genes that is particularly enriched in our dataset are ribosomal protein encoding transcripts that contain uORFs. While ribosomes were once thought to play a passive role in the selection of mRNA transcripts, recent studies suggest that specialized ribosomes contribute to the overall proteome through sequence-specific interactions (Xue and Barna 2012). The make-up of ribosomal proteins within a cell can influence the cell phenotypically, as evidenced by the tissue-specificity of certain ribosomal proteins (Kondrashov, Pusic et al. 2011, Guimaraes and Zavolan 2016). The relative contribution a 5’ TOP motif and translation of a uORF from the same 5’ leader in regulating downstream translation is unknown. It is possible that the 5’ TOP motif may exert similar regulation on the uORF as with the CDS, as we did observe that the majority of uORF/CDS pairs on these transcripts were repressed in neuronal differentiation. Interestingly, a uORF was detected on *RPS24* with a strong inverse relationship to the *RPS24* CDS. Mutations in *RPS24,* which lower the levels of this small ribosomal subunit protein, cause Diamond-Blackfan Anemia (DBA) (Gazda, Grabowska et al. 2006). Levels of this protein can affect the abundance of another key ribosomal protein mutated in DBA, *RPS19*, and has the potential to disturb the ribosomal stoichiometry necessary for proper translation (Gazda, Grabowska et al. 2006, Badhai, Frojmark et al. 2009, Shi, Fujii et al. 2017, Genuth and Barna 2018). uORFs are an interesting candidate for the regulation of this integral class of proteins.

In summary, we detected thousands of uORFs in a neuronal model, characterized their expression, and described an overarching role for uORFs as translational regulators during neuronal differentiation. Future studies will be needed to define how individual uORF/CDS relationships are regulated, but this work pinpoints neuronal differentiation as a cellular process that requires the function of uORFs to establish proper protein expression in these two cellular states.

## Methods

### SH-SY5Y cell maintenance and differentiation

SH-SY5Y cells were grown in DMEM:F12 media (Invitrogen) supplemented with 10% FBS, .01mg mL^-1^ Gentamicin and .25ug mL^-1^ Amphoreticin B. Cells were plated on 150mm plates that were either coated with .1mg/mL poly-D lysine (Millipore) for differentiation or uncoated. Cells were allowed to propagate to 80% confluency for 1-2 days prior to lysing for ribosome profiling or induction of differentiation. SH-SY5Y cells were differentiated for 6 days in 10μM retinoic acid (all-trans, Sigma), with media changed every 24 hours.

### Construction of the Ribosome Profiling libraries

Ribosome profiling libraries were prepared as in Ingolia et al., 2010 and Ingolia et al., 2012. Cells were washed with ice cold PBS with CHX at 100ug mL^-1^. Plates were immediately flash frozen in liquid nitrogen, moved to dry ice, and lysed (in the presence of CHX) to prevent ribosome loading and runoff. Additional lysates were processed in parallel for poly(A) mRNA purification and mRNA-sequencing library preparation. Polysomes were isolated from the ribosome footprinting lysates on a 1M sucrose cushion with high speed centrifugation using a 70.1Ti rotor (Beckman) at 55,000 r.p.m. for 4 hrs at 4°C. rRNA was eliminated prior to linker ligation using Ribo-Zero Gold rRNA Removal Kit (Illumina). Ribosome Profiling libraries were barcoded and multiplexed with 2-4 libraries per lane, and sequenced on a HiSeq 2000 (Illumina) using 50 cycles of single end reads. mRNA libraries were multiplexed on a single lane. All sequencing was conducted at the University of Michigan DNA Sequencing Core.

### Plasmid Construction

pcDNA 3.1 plasmid was modified to encode NanoLuc and GGG-NanoLuc as previously published (Kearse, Green et al. 2016). gBlocks^®^ (IDT) were ordered of the 5’ leader sequence to the last codon before the in-frame stop of selected genes flanked by restriction sites. These were restriction cloned upstream of GGG-nLuc using PacI and XhoI (NEB) with 12 nucleotides between the start of the 5’ leader and the T7 promoter sequence to reduce spurious initiation in sequences specific to the plasmid. Frameshifts were accomplished by PCR cloning with primers that inserted one or two nucleotides between the uORF and the nanoluciferase sequence. PCR products were cloned in place of nanoluciferase in the original uORF plasmid using XbaI and SacII (NEB). Restriction digest and Sanger Sequencing were used to confirm plasmid sequence.

### SH-SY5Y Transfection and Nanoluciferase Assay

SH-SY5Y cells were seeded on 6-well culture plates at 3 ×10^5^ cells per well. 24 hours post seeding, each well was transfected using 7.5μL FUGENE HD (Promega) and 1.25μg nanoluciferase reporter plasmid along with 1.25μg pGL4.13 (internal transfection control that encodes firefly luciferase [FFluc]) in 300 μL of OptiMEM (Invitrogen). Transfections of differentiated cells were performed on day 5 in RA supplemented media. Cultures were allowed to grow for 24 hours after transfection. Cells were lysed in 250μL Glo Lysis Buffer (Promega) for 5 minutes at room temperature. 50μL lysate was mixed with 50μL prepared Nano-GLO or ONE-Glo (Promega) for 2 minutes, and bioluminescence was detected using a GloMax^®^ 96 Microplate Luminometer. Nanoluciferase signal was normalized to FFluc signal in each sample. pcDNA 3.1 encoding nLuc the AUG start codon mutated to a GGG (GGG-nanoLuc) was run in parallel with each experimental nLuc plasmid and subjected to both conditions to serve as a control for normalization.

### Immunocytochemistry and microscopy

Cells were fixed at 37°C with 4% PFA/4% sucrose in PBS with 1mM MgCl2 and .1mM CaCl2 (PBS-MC), permeabilized for 5 minutes in .1% Triton-X in PBS-MC, and blocked for 1 hour with 5% bovine serum albumin in PBS-MC. Cells were incubated in blocking buffer and primary antibodies against β-actin (Santa Cruz Biotechnology, cat# sc-130656, 1:1000) and neurofilament (Abcam, Ab8135, 1:1000) for 1 hour at room temperature. Following 3×10 minute washes in PBS-MC, cells were incubated in PBS-MC with Alexa Flour 488 conjugated goat anti-rabbit IgG and Alexa Flour 635 conjugated goat anti-mouse IgG to achieve secondary detection (Thermo Fisher, 1:1000). Cells were washed again, and placed in ProLong^™^ Gold antifade reagent with DAPI (Invitrogen).

Imaging was performed on an inverted Olympus FV1000 laser-scanning confocal microscope using a 40x objective. Acquisition parameters were identical for each condition and optimized to eliminate signal bleed-through between channels. Images were converted to average-intensity z-projections in ImageJ. Cytoplasmic β-actin was quantified by averaging the integrated density corrected for background signal of the cells in each condition. The length of one main neurofilament-labeled primary neurite per cell was determined in ImageJ and converted from pixels to µm, and averaged for each condition.

### Western Blotting

Cells were maintained as described above. Cells were washed 2X in PBS, and RIPA buffer was added to a single well of a 12-well dish either at 80% confluency or after 6 days of retinoic acid differentiation. Cells were agitated for 40 minutes at 4°C to ensure complete lysis. Lysates were clarified by centrifugation, and the supernatant was mixed with reducing SDS sample buffer and boiled for 5 minutes at 90°C. Equal amounts of lysate were loaded on an SDS-PAGE gel and subsequent western blotting was carried out with primary antibodies against FMRP (1:1000, cat# 6B8, BioLegend), GAPDH (1:1000, cat# sc-32233, Santa Cruz Biotechnology), total eIF2α (1:1000, cat# 9722, Cell Signaling Technology), phospho-eIF2α (1:500, cat#: 44-728G, Invitrogen) or E7 Tubulin (1:1000, DSHB)—in 5% (wt/vol) non-fat dry milk in TBS-T (NFDM). An HRP conjugated goat antibody to mouse IgG or to rabbit IgG was used for secondary detection (1:5000, Jackson ImmunoResearch Laboratories) in 5% NFDM. SuperSignal^™^ West Femto Maximum Sensitivity Substrate (Thermo Scientific) was used for HRP detection of phospho-eIF2α levels. Western Lightening® Plus-ECL (PerkinElmer, Inc.) was used for all other antibody detection.

### Alignment to the human genome and transcriptome (GRCh38 Ensembl version 78)

Ribosome profiling and mRNA-Seq reads were trimmed of adapters, and then by quality using *fasqt-mcf* from the *ea-utils* package (Aronesty, 2011). Ribosome profiling and mRNA-Seq reads in FASTQ format were trimmed based on quality if four consecutive nucleotides were observed with Phred scores of 10 or below. The minimum read length required after trimming was 25 nucleotides.

Trimmed sequences were then aligned to a ribosomal RNA sequence index using Bowtie v1.1.2 (Langmead, *et. al.,* 2009) to deplete them of contaminant sequences. Alignment to the rRNA sequence contaminant index was performed using the following parameters: seed alignment length of 22 nucleotides, no mismatches in the seed alignment were allowed, with the unmapped reads written to a separate FASTQ file.

~~~
bowtie -l 22 -n 0 -S --un /path/to/depleted_reads.fq \
/path/to/rRNA_index \
/path/to/trimmed_reads.fq
~~~

Ribosome profiling and mRNA-Seq reads depleted of rRNA contaminant sequences were aligned to the human genome and transcriptome (Ensembl, version 78) using TopHat v.2.0.10 (Trapnell*, et. al.,* 2009). The trimmed and rRNA-deplete reads were aligned to the human genome and transcriptome with the parameters specifying standard Illumina reads, with un-gapped Bowtie 1 alignment (using a seed alignment length of 22 nucleotides, with no mismatches in the seed alignment allowed), to annotated junctions only, using Solexa quality scores:

~~~
tophat2 -p 4 –bowtie1 \
–-no-novel-juncs \
--library-type fr-unstranded \
--solexa-quals \
-G /path/to/ensemble.gtf \
/path/to/bowtie_index \
/path/to/depleted_reads.fq
~~~

### Sequence alignment quality filtering and merging

Ribosome profiling and mRNA-Seq reads aligned to the human genome and transcriptome by TopHat2 were output to BAM format, and then sorted by chromosomal coordinate. Reads were then filtered by mapping quality using SAMtools (Li, *et. al.,* 2009); read alignments were required to have minimum mapping quality of 10, or higher, to be retained for subsequent analyses. Unique read group identifiers were assigned to each technical and biological replicate, and then the alignments were merged by technical replicates and subsequently as biological replicates using Picard (http://broadinstitute.github.io/picard/).

### Metagene profile generation and alignment offset calculation

For counting reads over transcript isoforms, metagene profiles were generated from the Ensembl (version 78) transcript annotation database using Plastid (Dunn*, et. al.*, 2016). A- and P-site offsets for harringtonine and cycloheximide ribosome profiling reads, respectively, were determined by pooling all reads that overlapped canonical AUG translation initiation start sites from annotated protein-coding genes. The most common (mode) distance from the 5’ ends of reads of a given length to the position of the AUG in those reads was accepted as the A- or P-site offset distance.

### Calculation of transcript abundance

Read counts over each transcript isoform, or region (5’UTR, CDS, and 3’UTR), were normalized by length, summed, and reported as transcripts per million (TPM) as described previously (Li, *et. al.,* 2011). At the time of analysis, Cufflinks (Trapnell, *et. al.,* 2010) was required for initial transcript quality control, and was run with the following parameters:

~~~
cufflinks -p 8 -o /path/to/output \
-G /path/to/ensemble.gtf \
/path/to/tophat/alignments
~~~

### SPECtre analysis of transcripts in non-differentiated and RA-differentiated libraries

SPECtre profiling (Chun, *et. al.,* 2016) measures the strength of the tri-nucleotide periodicity inherent to the alignment of ribosome protected fragments to protein-coding genes in a reference transcriptome. SPECtre analysis was applied in two stages: 1) to score the translational potential of annotated transcripts to build a background protein-coding reference model, and 2) to score the translational potential of predicted upstream-initiation ORFs. In this way, the translational potential of predicted upstream and overlapping ORFs are score against a background model of translation derived from annotated protein-coding transcripts. Annotated protein-coding transcripts were profiled by SPECtre using the following parameters:

~~~
python /path/to/SPECtre.py \
--input /path/to/tophat/alignments \
--output /path/to/output \
--log /path/to/logfile \
--gtf /path/to/ensemble.gtf \
--fpkm /path/to/cufflinks/isoforms.fpkm_tracking \
--len 30 \
--fdr 0.05 \
--min 3.0 \
--nt 8 \
--type mean \
--target <chromosome_id>
~~~

Where the minimum FPKM required for a transcript to be considered as translated for generation of the background model was specified as 3.0, and the length of the sliding SPECtre windows was set to encompass 30 nucleotides. The SPECtre score for a transcripts was defined as the mean of the scores over these sliding windows, and a 5% false discovery rate was established to set the minimum SPECtre translational score threshold. In addition, SPECtre profiling was split by chromosome to speed computation, and the results were merged afterwards using a custom Python script. Finally, prior to analysis of predicted upstream-initiated ORFs by SPECtre profiling, the minimum SPECtre translational threshold was re-calculated using TPM instead of FPKM using a minimum cutoff of 10 transcripts per million.

### Computational prediction of upstream-initiated open reading frames

Open reading frames were computationally predicted from annotated 5’UTR sequences (Ensembl, version 78) using AUG, and near-cognate non-AUG translation initiation site sequences. Open reading frame sequences were generated based on these predicted initiation site sequences and read through to the first in-frame termination codon encountered in the annotated CDS. These predicted ORFs were then used to generate coordinates over which they would be profiled and scored by SPECtre. Identical parameters to the annotated transcript SPECtre analysis were employed for consistency across analyses:

~~~
python /path/to/SPECtre-uORFs.py \
--input /path/to/alignments \
--output /path/to/output \
--results /path/to/spectre/transcript_results \
--log /path/to/logfile \
--fpkm /path/to/cufflinks/isoforms.fpkm_tracking \
--len 30 \
--fdr 0.05 \
--min 3.0 \
--nt 8 \
--type mean \
--target <chromosome_id>
~~~

Three alternative inputs are required for the SPECtre analysis of predicted ORFs: 1) the annotated transcript GTF database was not required and removed, 2) the results of the annotated transcript analysis, and 3) a genomic sequence file in FASTA format. The results of the annotated transcript analysis were used to identify the set of transcripts from which to predict upstream-initiated ORFs, and the FASTA sequence file was used to generated the ORF sequences for output.

### Supplemental annotation of non-AUG translation initiation sites

Upstream sequences of predicted non-AUG translation initiation sites were examined for possible in-frame AUG initiation start sites; 5’UTR sequences of predicted non-AUG sites were extracted, and then searched for the presence of in-frame AUG sites. These non-AUG sites were then re-annotated according to the proximity of upstream AUG initiation sites: those with an in-frame AUG site within 30 nucleotides of the predicted TIS, and those with an in-frame AUG site in-frame, but beyond 30 nucleotides upstream of the predicted site.

### Kernel density estimation of differential uORF translation on CDS translational efficiency

To further differentiate those uORFs with differential translation and identify those that contribute to the regulation of downstream CDS, the log-change in predicted ORF TPM was compared against the log-change in downstream CDS TPM across the conditions. The differential translational identity of each predicted ORF was retained from the SPECtre clustering analysis, and kernel density estimation was performed using R.

### Heuristic filtering of predicted uORFs

Candidate uORFs were filtered based on a series of heuristic criteria, including: 1) the removal of predicted uORFs with no RPF coverage in the 5’UTR of the transcript, 2) the removal of uORFs predicted to initiate within 15-nt of the annotated CDS start site, and 3) the removal of predicted uORFs without matching mRNA-Seq coverage in the 5’UTR of the transcript. Following these initial minimal coverage filters, the candidate uORFs were further stratified by quality of coverage. First, identical uORF isoforms in overlapping transcripts within the same protein-coding gene annotation set were merged into a single candidate. Next, overlapping uORF candidates were prioritized by the extent of RPF coverage to their 5’ end, as well as by overall coverage. Finally, any remaining overlapping uORF candidates were prioritized by the magnitude of their calculated SPECtre score, with higher scored candidates preferred. R code for the functions to execute the heuristic filtering of uORF candidates is replicated in **Supplemental Methods**.

### Calculation of translational efficiency

Ribosome profiling or mRNA-Seq reads were counted over each region (5’UTR, CDS, and 3’UTR), transcript, or uORF and then normalized to length and library size as transcripts per million (Li and Dewey 2011).(Li and Dewey 2011). To calculate translational efficiency over a region, transcript or uORF, ribosome profiling TPM in each biological replicate across each condition was quantile normalized (Amaratunga*, et. al.,* 2001) and then divided by the quantile normalized TPM in mRNA-Seq. Read and RPF counts from mRNA-Seq and ribosome profiling libraries does not include those that overlap the 5’UTR and 3’UTR. Furthermore, to limit the boundary effects due to translation initiation and termination, RPF and read counts do not include those reads whose A- or P-site adjusted position for harringtonine and cycloheximide libraries, respectively, overlap the first or last 15 nucleotides of an annotated CDS.

### Differential expression analysis and gene set enrichment testing in mRNA-Seq

As described previously, the read abundance over annotated protein-coding transcripts was calculated as TPM, then quantile normalized across conditions using the preprocessCore package (Bolstad, 2016) in R (R Core Team, 2017), and then ranked. The change in rank for each gene was calculated across the non-differentiated and RA-differentiated conditions, and the significance of the up- or down-regulation of these rang-changes across conditions was classified using an extreme outlier cutoff (Tukey 1949).(Tukey 1949). Functional characterization of these significantly rank-changed genes across the non-differentiated and RA-differentiated conditions was analyzed using the goseq package (Young, Wakefield et al. 2010) in R, and corrected for multiple testing using Benjami-Hochberg adjusted p-values.

### Differential translation analysis and gene set enrichment testing in Ribosome Profiling

Ribosome profiling read fragments were A- or P-site adjusted, and then counted over annotated protein-coding CDS regions in each biological replicate using the metagene profiles generated by Plastid (Dunn and Weissman 2016). As described previously, ribosome-protected fragments with A- or P-site adjusted positions that overlapped the first or last 15 nucleotides of the boundaries defined by the annotated CDS region were masked from the analysis. DESeq2 (Love, Huber et al. 2014) was used to identify those genes with differential translation across the two states of cellular differentiation. Genes were annotated as significantly up- or down-regulated using a Benjamini-Hochberg adjusted p-value cutoff of 0.1, and fold-change in counts greater than 1, or less than 1, respectively. Functional characterization of these significantly up- and down-regulated genes was analyzed by goseq using parameters specified previously.

### Differential translational efficiency and gene set enrichment testing in Ribosome Profiling

For each biological replicate, ribosome profiling read fragments were A- or P-site adjusted, and then counted over annotated protein-coding CDS regions using the metagene profiles generated by Plastid. As above, read counts over the first and last 15 nucleotides of protein-coding CDS regions were masked for subsequent analyses. In addition, mRNA-Seq read counts were extracted from each condition, with the proximal and terminal 15 nucleotide ends of the CDS masked for consistency with the RPF counts. The DESeq2 wrapper for differential translational efficiency analysis, Riborex (Li, Wang et al. 2017), was used to identify those genes with significant changes in translational efficiency. Genes were annotated as significantly up- or down- regulated using a Benjamini-Hochberg adjusted p-value cutoff of 0.1, and absolute fold-change of 1. Functional characterization of the sets of genes enriched in each condition by translational efficiency was analyzed by goseq using parameters described previously.

### Conservation analysis

To assess the conservation of the various regions, transcripts and uORFs, the PhyloCSF scores (Lin, Jungreis et al. 2011) over each target region was extracted. For uORFs, the PhyloCSF score was extracted according to its predicted phase. In order to de-convolute the contribution of regional conservation due to overlap with annotated CDS regions, predicted uORFs that did not initiate and terminate wholly upstream of a CDS were also scored according to the subset of their coordinates defined by the 5’UTR alone. The mean PhyloCSF over each of these regions and uORFs was calculated, and then mean-shifted to the canonical (+0) reading frame of the annotated CDS for comparison.

### GC nucleotide content analysis

Similar to the conservation scores, the ratio of GC nucleotide content in each reading frame of 5’UTRs, and CDS. GC content over the predicted phase of each uORF was calculated, with the 5’UTR overlapping region of CDS-terminated uORFs deconvoluted from the region overlapping the CDS as described above.

### Cluster analysis of differential uORF translation by SPECtre score

In order to identify subsets of uORFs with differential translation in one state of cell differentiation compared to the other, the SPECtre score for each predicted uORF was calculated (described in Supplemental Materials and Methods). The SPECtre score of each predicted uORF was classified by k-means clustering in R to define sets of uORFs with differential translation in one of the conditions, and those with no difference in translational potential between the two conditions.

### Additional filtering of candidate uORFs

Additional replicate-based filtering was applied to the set of predicted uORFs to identify a set of highly confident candidates. As a form of internal validation, a predicted uORF was required to meet a minimal translational threshold in at least one of the biological replicate samples across both conditions. This threshold was determined on a conditional basis dependent on the 5% FDR cutoff required for translational activity according to the distribution of SPECtre scores in protein-coding genes.

## DATA AVAILABILITY

The datasets generated for this study will be posted to the GEO database upon formal publication.

## AUTHOR CONTRIBUTIONS

PKT, CMR and REM conceived the project. CMR conducted all of the RP experiments molecular and cellular biological studies. SYC performed all of the bioinformatics analysis with guidance from REM. All authors analyzed and guided the data analysis strategy. CMR and PKT wrote the manuscript with significant input from SYC and REM. All authors edited the manuscript.

## ACKNOWLEDGEMENTS

We would like to thank the University of Michigan DNA Sequencing Core for their advice, technical support, and services.

